# Natural Amelioration of Mn-induced Chlorosis Facilitated by Mn Down-regulation, Ammonium and Rainwater in Sugarcane Seedlings

**DOI:** 10.1101/618124

**Authors:** Gui Zhi Ling, Xiao Xia Wang, Shu Yang, Xin Lian Tang, Shi Jin Jia, Min Min Chang, Xiao Feng Li

## Abstract

We had previously reported that manganese (Mn)-induced chlorosis is a serious problem in ratoon sugarcane seedlings grown in acidic soils. To further monitor the progression of chlorosis and elucidate the corresponding mechanism, both plant growth and nutrient status of sugarcane plants and soils were investigated in the growth seasons of ratoon cane seedlings in 2016 and 2018. The impacts of rainfall and ammonium on chlorosis were also investigated hydroponically. The results showed that the chlorotic seedlings could green in mid-summer; Mn content in the first expanded leaf decreased significantly, whereas iron (Fe) content increased significantly during the progression of greening. The leaf Mn content in the greened seedlings decreased by up to 78.1% when compared with that in the initial chlorotic seedlings. The seedlings also showed a significant increase in seedling height and weight of the expanded leaves, accompanied by a decrease in plant Mn content during the progression of greening. Moreover, young seedlings with less Mn content showed earlier greening than older seedlings with more Mn content. The exchangeable ammonium content in the soils increased significantly during the progression of greening, and the addition of 1 mM ammonium to the chlorotic seedlings resulted in a decrease in leaf Mn content by up to 80%. Furthermore, leaf SPAD value and Fe content increased by 2.0-fold and 1.4-fold, respectively, after rainwater was applied to the chlorotic seedling. These results indicate Mn-induced chlorotic seedlings can turn naturally green, and downregulation of plant Mn content, rainfall in summer, and soil ammonium contribute to the greening of chlorotic seedlings.

## Introduction

Manganese (Mn) is an essential element for normal plant growth and development, but it is toxic when present in excess. Mn toxicity is a major limiting factor for crop growth and yield in acidic soils, which contains an abundance of Mn^2+^ [1]. Because plant roots absorb Mn as a divalent cation, any factors that change its concentration in the soil solution can potentially affect the accumulation of Mn in the plant [2]. The incidence of Mn toxicity strongly depends on the status of nutrients in the growth medium and concentrations of other elements that affect Mn absorption, translocation, and use [2]. Most of the studies on the interaction between iron (Fe) and Mn in plants have reported a negative correlation between Fe and Mn accumulation in the plant shoots [2]. Fe supply has a marked effect on alleviating Mn toxicity in many crops. The form of nitrogen (N) also affects Mn toxicity [3]; rice plants grown with ammonium have less Mn in their shoot tissue and develop slight or no symptoms of Mn toxicity. Mn transporters are involved in the regulation of nitrate-induced Mn accumulation in rice [4]. The incidence of Mn toxicity also strongly depends on the environmental factors. High temperature reduces the growth rate of and increases Mn absorption by plants [5]. High light intensity has been reported to stimulate Mn absorption by plants and accentuate the severity of Mn toxicity [6].

Sugarcane (*Saccharum officinarum* L.) is an important crop for sugar production worldwide. Recently, Huang et al., authors in our laboratory, reported Mn-induced chlorosis in ratoon sugarcane grown in acid soils [7]. However, the progression of chlorosis and involvement of soil and environmental factors in this progression are still unclear. It is important to analyze the causes of chlorotic seedling greening and elucidate corresponding mechanisms for developing problem-solving strategies in agricultural practices. Therefore, in the present study, we monitored the development of chlorosis and investigated sugarcane Mn and Fe contents and soil ammonium content during the progression of chlorotic seedling greening in the growth seasons of 2016 and 2018. To evaluate the significance of rainfall and ammonium and impacts of rainwater and ammonium on the greening, Mn and/or Fe contents in sugarcane were also studied using hydroponic experiments.

## Materials and methods

### Monitoring Mn-induced chlorosis in ratoon cane seedlings

This study was initiated on April 15 (day 0), 2016 and 2018, in Chongzuo, Guangxi Province, China, where Mn-induced chlorosis in ratoon cane has been widely found in these past few decades [7]. This region has a typical subtropical monsoon climate, with an annual precipitation of 1200 mm. Two fields in the same area were selected to investigate greening of the chlorotic seedlings and changes in the Mn and Fe contents of the plants and ammonium content in the soils as the greening progressed. In the first field, chlorotic seedlings, young plants (YP), bourgeoned in April, whereas in the second field, chlorotic seedlings, older plants (OP), bourgeoned in January. The soils were strongly acidic, with 2.5, 5.8 mg·kg^−1^ soluble Mn and 4.0 pH for YP and OP soils, respectively.

The extent of chlorosis and greening was investigated on day 0, 5, 10, 25, 35, 45, and 60 in 2016 and day 0, 10, 40, and 58 in 2018. Simultaneously, seedling height and leaf SPAD values were measured, and the leaves and soils were sampled. The chlorotic and greened leaves of the YP seedlings were recorded with a camera (Canon, EOS 5D Mark IV, Japan) on day 0 and 40, respectively.

The seedling height and SPAD values of the first expanded leaf were measured with a ruler and chlorophyll meter (SPAD-502Plus; Minolta, Japan), respectively. Ten seedlings were measured, and the same seedlings were measured at different time points. To investigate changes in leaf weight during the progression of greening, overall expanded leaves were collected from 10 seedlings on each site at each time point and weighed. The first expanded leaf was collected randomly to determine the Mn and Fe contents, and 30 leaves were included in each sample. Separation of the cell sap of the first expand leaf was performed using the method described by Wang et al. [8]. The Mn content in the leaves was analyzed with an atomic absorption spectrometer (PINAACLE900T; PerkinElmer, USA), after the leaves were cleaned, dried, and crushed according to the method published by Huang et al. [7]. Fe content in the leaves was measured spectrophotometrically (Shimadzu UV-1800; Kyoto, Japan), according to the method reported by Mehrota et al. [9], after the leaves were cleaned and cut into 0.5 mm segments with stainless steel scissors.

Soil samples were collected from 0–20 cm of the topsoil layer. Exchangeable ammonium in these samples was determined with a continuous flow analytical system (SEAL-AA3, SEAL, Germany), after extraction with KCl by using the method reported by Ruan et al. [10].

### Culture experiment

A sugarcane cultivar, G32, was used in hydroponic experiments. The sugarcane seedlings were prepared and cultured according the methods described by Yang et al. [11]. The seedlings were cultured in a growth chamber with a controlled environment: 14 h (30 °C)/10 h (25 °C) day/night cycle, relative humidity of 75–85%, and daytime light intensity of 400 μmol m^−2^ s^−1^.

To investigate the impact of ammonium on Mn content in the plants, 15-day-old seedlings were exposed to aerated nutrient solution (pH 5.5) with 0.5 mM MnCl_2_ and 0 or 1.0 mM NH_4_Cl (ammonium). The base nutrient solution was nitrogen-free 1/5 strength Hoagland solution. The first expanded leaf was collected after the treatment with ammonium for 0, 4, 8, and 12 days. The leaf Mn content was determined as described above.

To prepare Mn-induced chlorotic ratoon cane seedlings in a controlled environment, the plants were exposed to 1/5 strength Hoagland solution containing 0.5 mM MnCl_2_ for 30 days. Then, the shoots were excised from the stem bases, and the roots were exposed to 0.5 mM CaCl_2_ solution. After the new-generation seedlings were bourgeoned from Mn pre-cultured plants, the roots of the seedlings were exposed to 1/5 strength Hoagland solution containing free Mn and Fe. Fifteen-day-old ratoon cane seedlings showing visible symptoms of Mn-induced chlorosis were treated with rainwater (+RW) or not treated (-RW) to investigate the effects of rainwater on chlorosis. The rainwater was collected with a plastic container during rainfall in May 2018. Fe concentration in the collected rainwater was 0.26 mg·L^−1^. The rainwater treatment was performed by exposing the roots to aerated rainwater and spraying the leaves of the seedlings with rainwater three times a day: morning, afternoon, and evening. Deionized water, instead of rainwater, was used for the leaves and roots of the control ratoon cane seedlings (-RW). After 15 days, SPAD values of the first expanded leaves were measured, and the plants were harvested. The first expanded leaves of the seedlings treated with –RW or +RW were photographed with the camera. The contents of Mn and Fe in the leaves were determined as described above.

### Statistical analysis

The data were analyzed by analysis of variance, and the treatment means were compared by Duncan’s multiplerange test or Student’s t test.

### Ethics statement

All experiments carried out in this study comply with current laws of China. Soil and plant samples were collected with oral permission from farmers to avoid interference with normal agriculture activities. The studies did not involve endangered or protected species.

## Results

### Reversal of Mn-induced chlorosis in the sugarcane seedlings

Excessive Mn-induced chlorosis in the sugarcane seedlings grown in acidic soils was monitored in 2016 and 2018. The seedlings exhibited visible symptoms of chlorosis in late spring (Fig 1A), extremely low levels of chlorophyll in the leaves, and SPAD values as low as 10 (Fig 2). However, the chlorotic leaves turned naturally green in early or mid-summer (Fig 2B). Although the chlorophyll contents on day 5, 10, and 25 in 2016 (A) and day 10 in 2018 (B) were similar to those on day 0, the contents significantly increased thereafter. On day 58 and 60 for OP seedlings and day 40 and 45 for YP seedlings, when the leaf SPAD values were greater than approximately 45, the visible symptoms of chlorosis disappeared. These results indicate that Mn-induced chlorosis can be naturally alleviated with season alternation.

**Fig 1.**
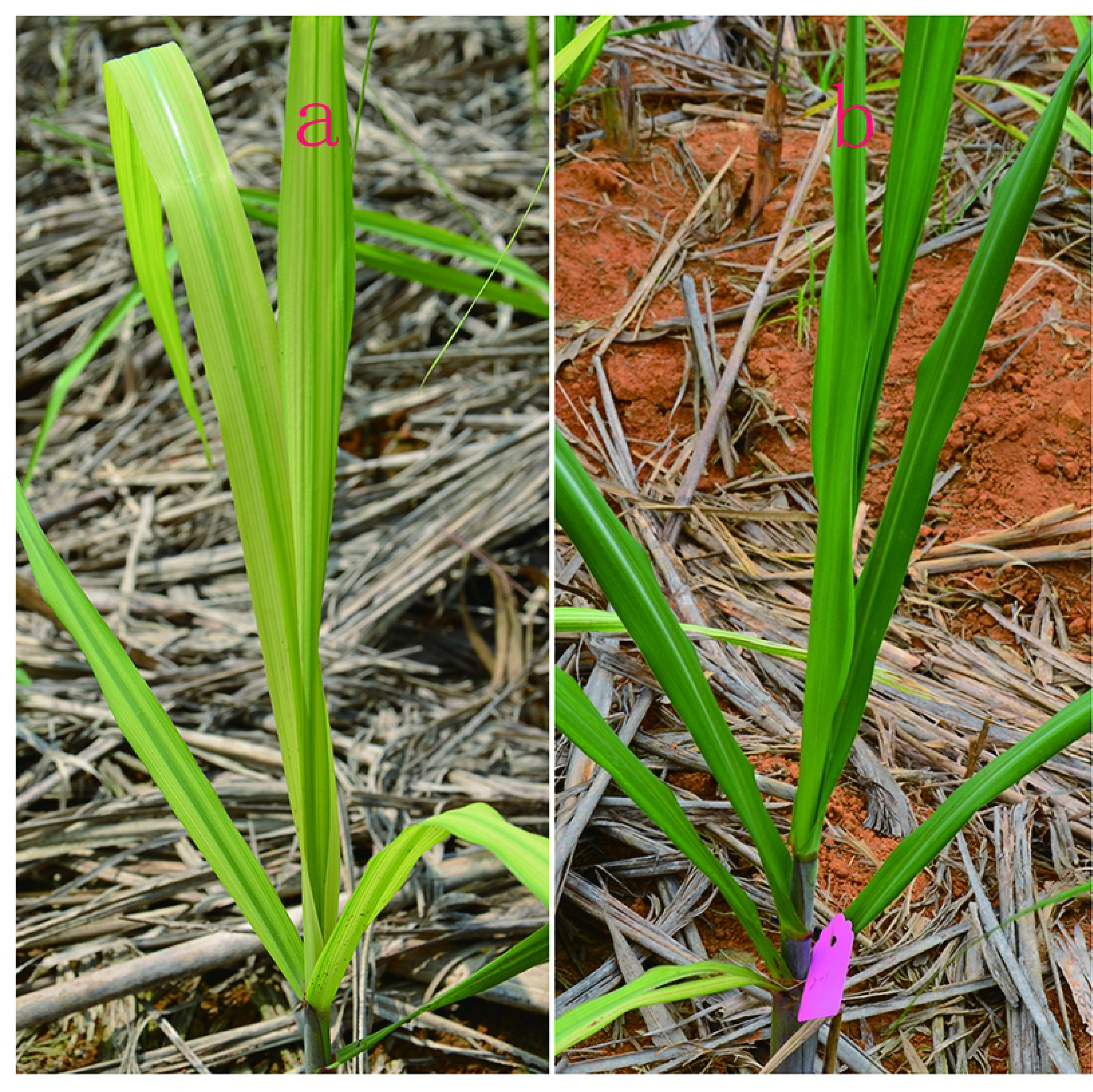
Etiolated (a) and greened (b) seedlings grown in an acidic soil.

**Fig 2.**
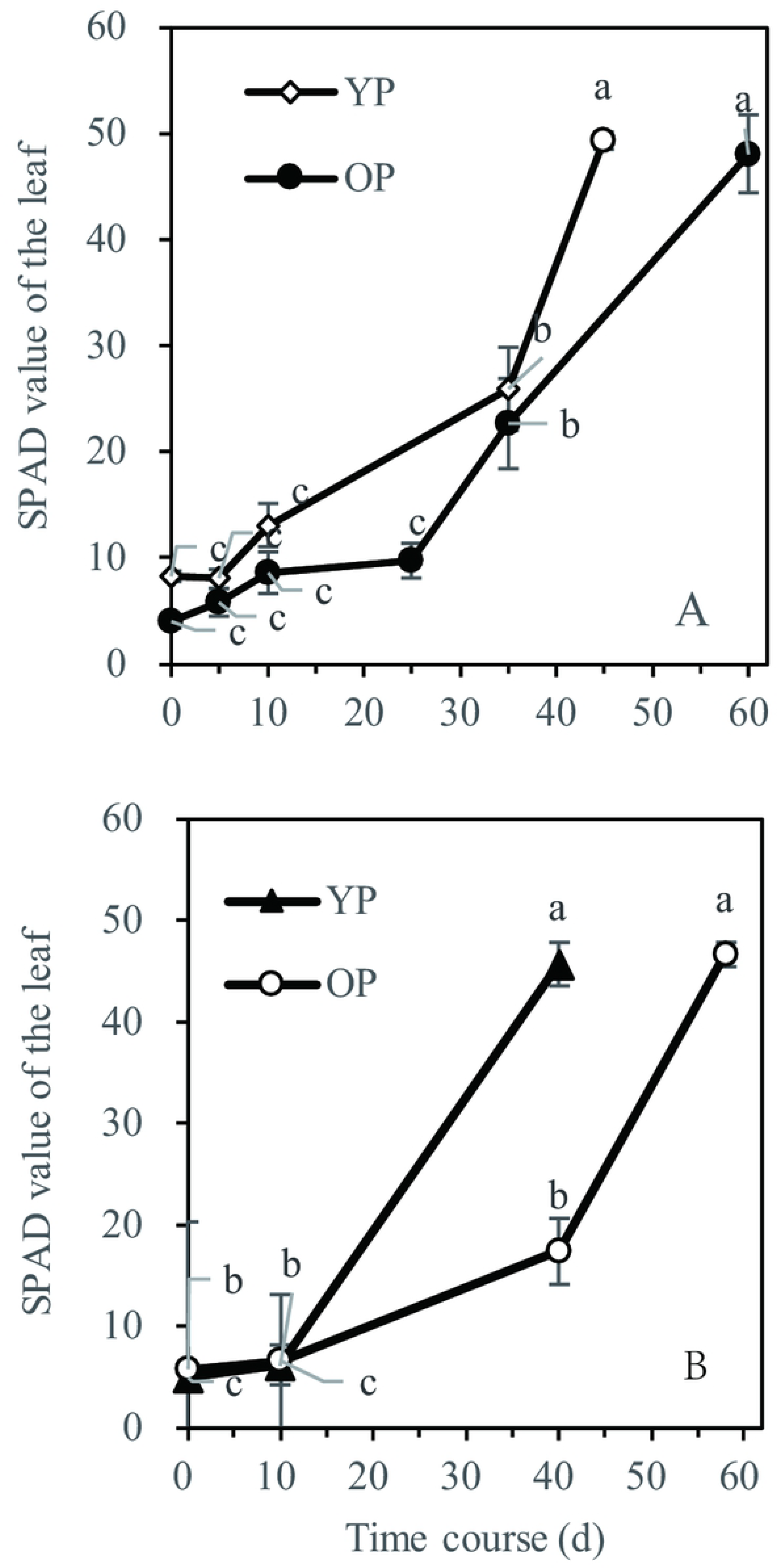
Changes in chlorophyll concentration, as SPAD value, in the first expanded leaf of the seedlings during the progression of greening. The values were first measured at April 15, 2016 (A) and 2018 (B) (day 0) and measured on the indicated days after the first measurement. YS, seedlings that bourgeoned in April; OS, seedlings that bourgeoned in January. Each value is the mean ± SEM (n = 3 replicates). Different letters on the same line indicate that the values are significantly different at the 0.05 level, according to Duncan’s multiple range test, the same as the below.

The greening was associated with seedling age. Visible symptoms of chlorosis were not observed on May 25 (day 40), 2018, in the YP seedlings, and the leaf SPAD value was 45.7; however, the symptoms were observed in the OP seedlings, which showed a lower SPAD value (17.5; Fig 2B). When the tracking period extended to 60 days, the visible symptoms of chlorosis disappeared in the OP seedlings as well, and the SPAD value increased to 46.7. A trend of early greening was observed in the YP seedlings grown in 2016 (Fig 2A).

### Mn content in the seedlings

The Mn content in the plants significantly decreased as the greening progressed (Fig 3A and B). The Mn content in the leaf was greater than 500 mg·kg^−1^ DW on day 0, and it decreased to 153.4–210.6 mg·kg^−1^ DW in the green plants on day 40–60; the rate of decrease ranged from 62% to 78.1%. The leaves of the chlorotic OP seedlings contained 266.1 mg·kg^−1^ DW Mn on day 35, whereas the Mn content in the other chlorotic seedlings was higher than 266.1 mg·kg^−1^ DW (Fig 3A). On day 40, the OP seedling with 342.5 mg·kg^−1^ DW Mn in their leaves showed visible symptoms of chlorosis; in contrast, the Mn content in the YP seedlings was less than 250 mg·kg^−1^ DW (243.5 mg·kg^−1^ DW), and the plants showed visible greening (Fig 3B). Mn in the leaf cell sap of the initially chlorotic seedlings and greened seedlings are shown in Fig 3C. The distribution of Mn to leaf cell sap decreased significantly after the chlorotic seedlings greened. These results indicate that the greening of chlorotic seedlings is a result of the decrease in Mn content in the plant, rather than Mn accumulation in the cell sap.

**Fig 3.**
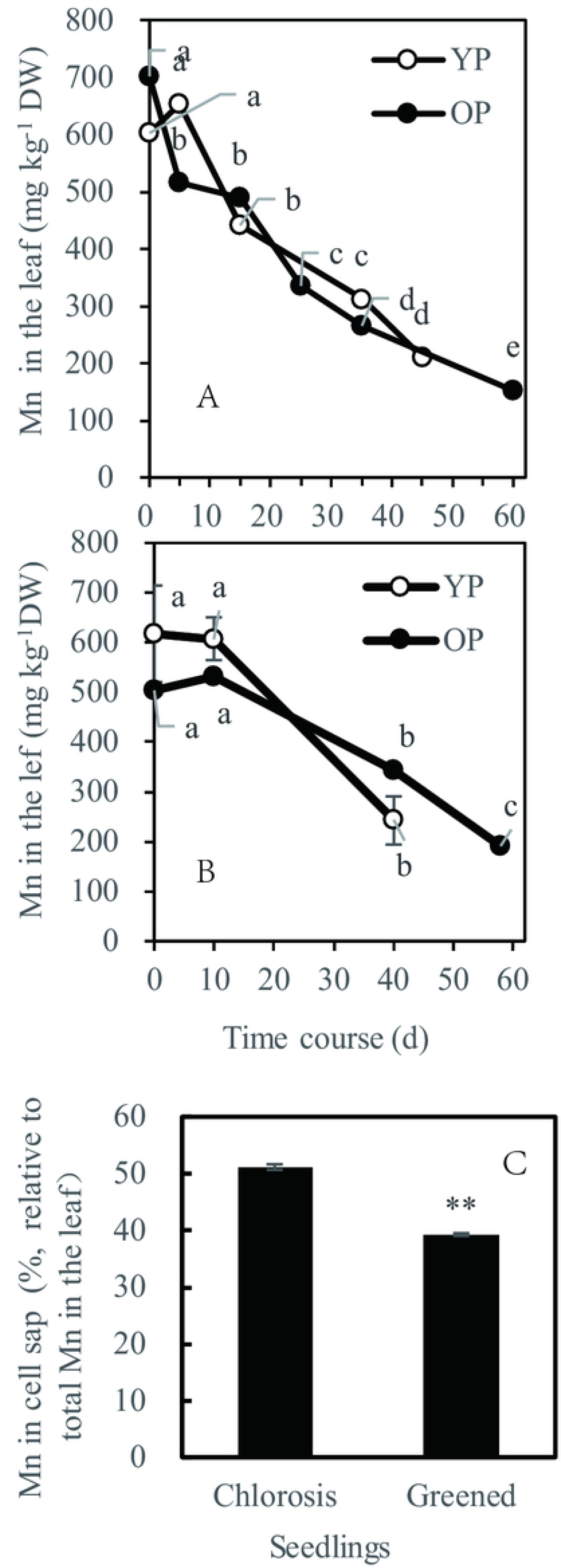
Changes in leaf Mn with the progression of greening of the chlorotic seedlings. The first expanded leaf was collected on the indicated days in 2016 (A) and 2018 (B). Mn distribution to cell sap of the first expanded leaf (C).

### Seedling growth

Natural greening of chlorotic seedlings was accompanied by the improvement of plant growth (Fig 4). No significant differences in leaf weight and seedling height were observed between the plants on day 0 and 10. However, a significant increase in leaf weight and seedling height was detected on day 40 (Fig 4), when the leaf SPAD value increased significantly (Fig 2). The changes in leaf weight and seedling height during the progression of greening were the opposite of those in leaf Mn content, which decreased as the greening progressed (Fig 3A). The soils showed no significant variations in both available and soluble Mn contents during the study (data not shown). These results imply that plant growth could be a reason for the decrease in plant Mn content.

**Fig 4.**
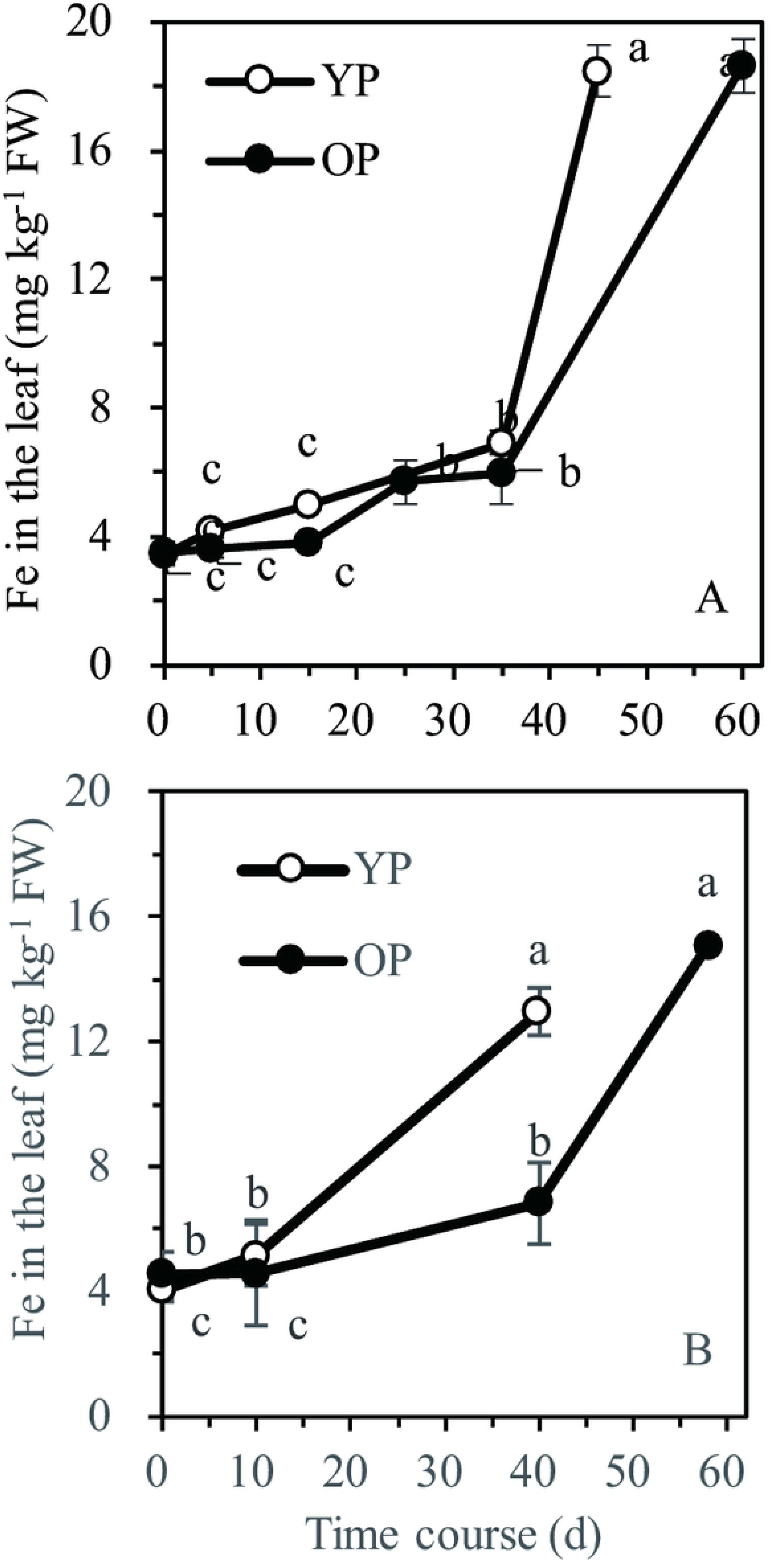
Changes in leaf Fe content with the progression of greening. The first expanded leaf was collected on the indicated days in 2016 (A) and 2018 (B).

### Fe content in the seedlings and impact of rainwater on chlorosis

The Fe content in the seedlings increased significantly as the greening progressed, and the greened seedlings showed higher Fe content in their leaves (Fig 5). All the chlorotic seedlings contained less than 10 mg·kg^−1^ FW of Fe in the leaves. On day 40, 45, 58, and 60, Fe contents in the leaves of the greened seedlings reached 12.9, 18.5, 15.1, and 18.7 mg·kg^−1^ FW, respectively. However, the soils showed no significant differences in exchangeable and soluble Fe contents during the study (data not shown).

**Fig 5.**
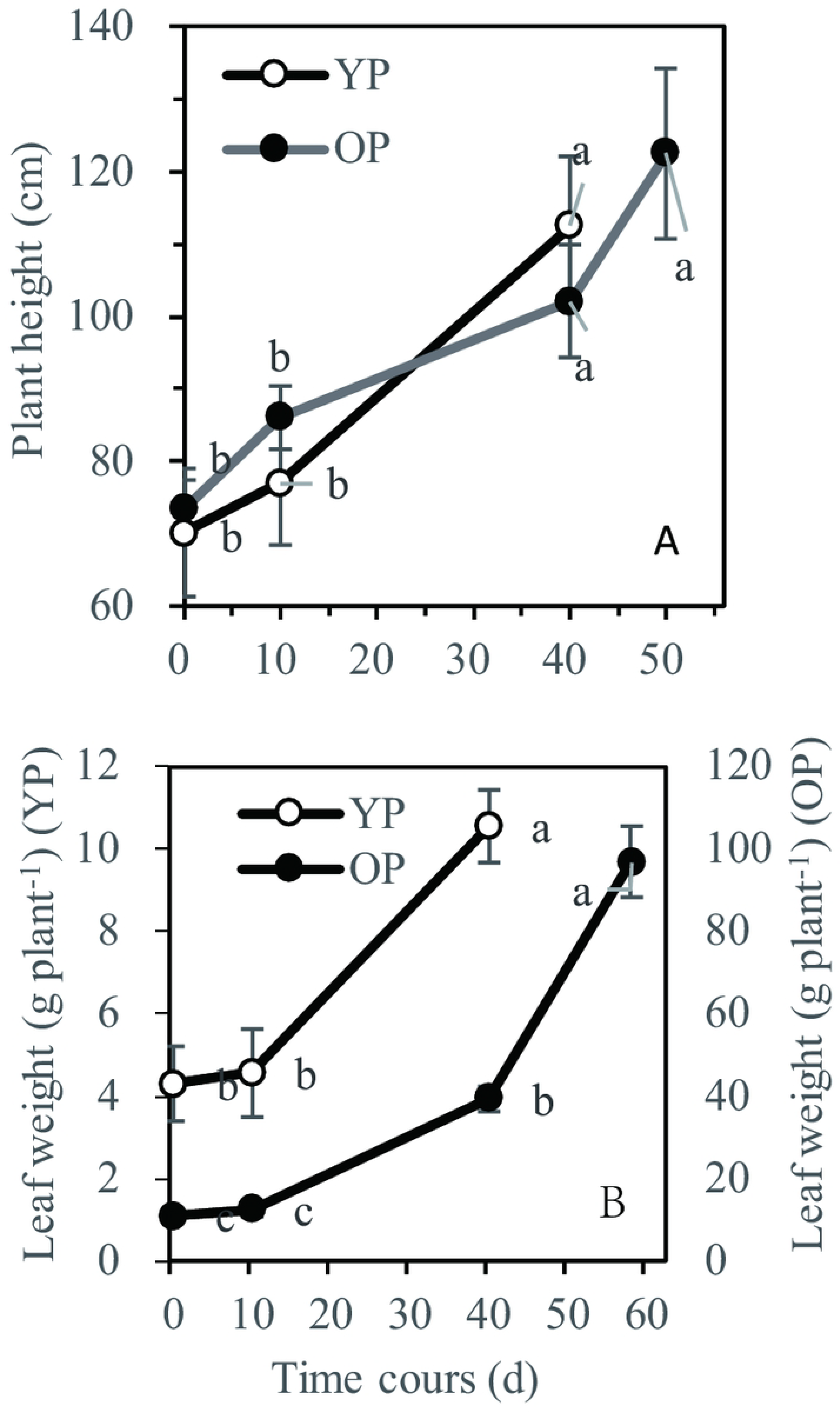
Changes in seedling height and total weight of the expand leaf blades during the progression of greening. The blades were collected from the seedlings on the indicated days in 2018.

Rainy season in the study area begins in early-summer, and the precipitation is extremely high (Fig S). To determine the significance of rainwater for the greening of the seedlings, effects of rainwater on the chlorotic ratoon seedlings were investigated (Fig 6). The symptoms of chlorosis were effectively alleviated, and the chlorotic leaves obviously greened after the application of rainwater for 15 days (Fig 6A). The leaf SPAD value of the control plants (-RW) did not vary significantly during the 15 day period of the treatment, whereas the value increased significantly (2-fold), after the application of rainwater (+RW; Fig 6B). The rainwater treatment did not influence Mn content in the leaves (Fig 6C). However, Fe content in the leaves increased significantly (1.4-fold) after the rainwater treatment. These results suggest that Mn-induced chlorosis was alleviated by an increase in plant Fe content due to rainwater.

**Fig 6.**
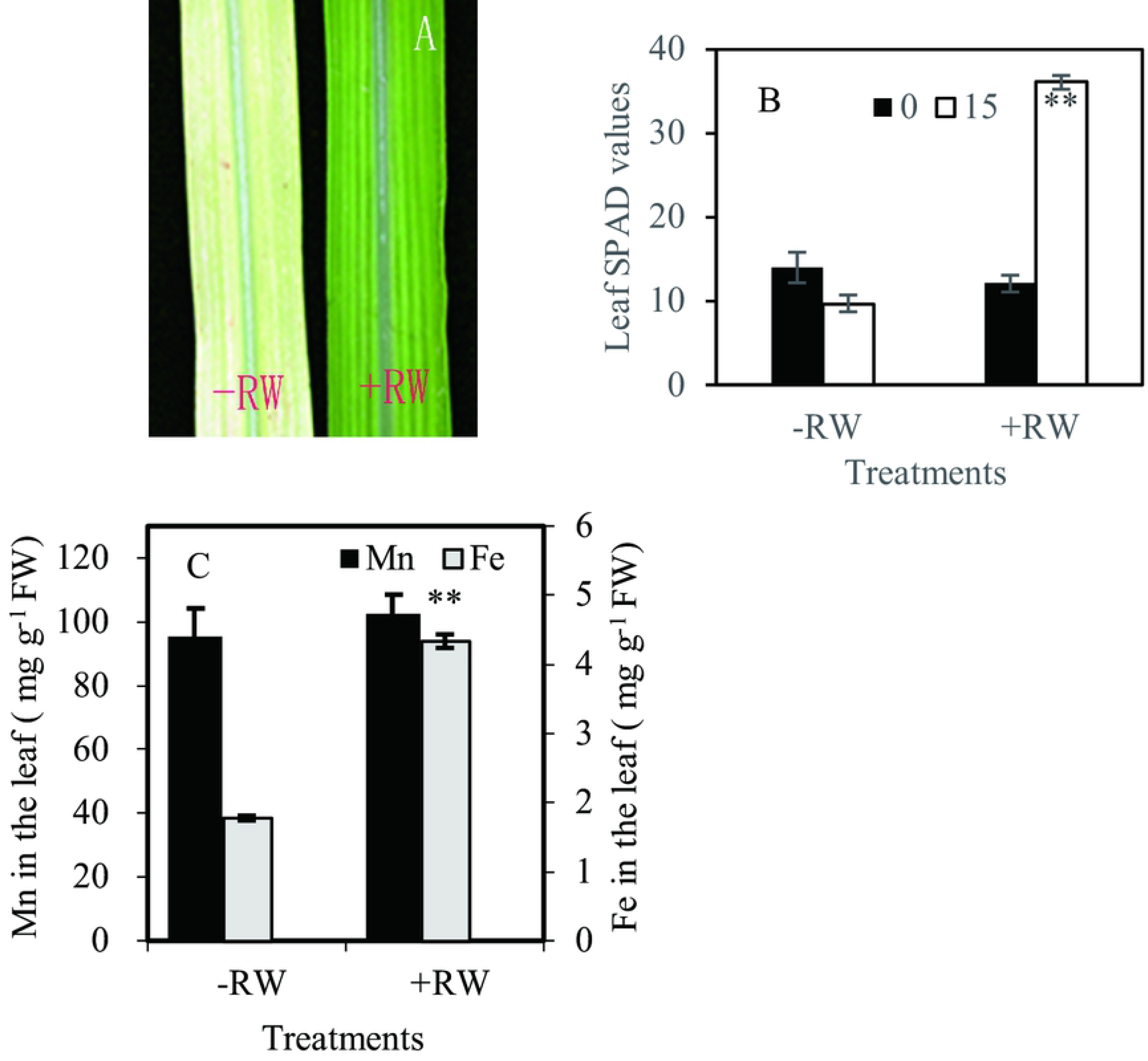
Effects of rainwater on the greening of Mn-induced chlorotic seedlings. Leaves of ratoon cane seedlings were exposed to rainwater (+RW) or not exposed (-RW), with no Mn addition for 15 days (A). Chlorophyll (B) and Mn and Fe (C) contents in the first expanded leaf after the seedling roots were exposed or not exposed to rainwater for 0 or 15 days. The ratoon cane seedlings bourgeoned from plants of the previous growth season exposed to a nutrient solution with 0.5 mM MnCl_2_ solution for 30 days. ** on the same parameter indicates significant difference at the 0.01 level, according to the pairwise Student’s *t*-test.

### Ammonium in the soils and effects of ammonium on Mn accumulation

The exchangeable ammonium content in the soils increased significantly as the greening progressed (Fig 7). Although the ammonium content was similar in the same soil on day 0 and 10, it increased significantly thereafter. The greening YP seedlings showed a sharp increase in exchangeable ammonium after 10 days, whereas the late-greening OP seedling showed a moderate increase. When the seedlings greened on day 40 and 58, the ammonium contents in the OP and YP seedlings increased by 3.7- and 6.2-fold, respectively, when compared with the ammonium content on day 0.

**Fig 7.**
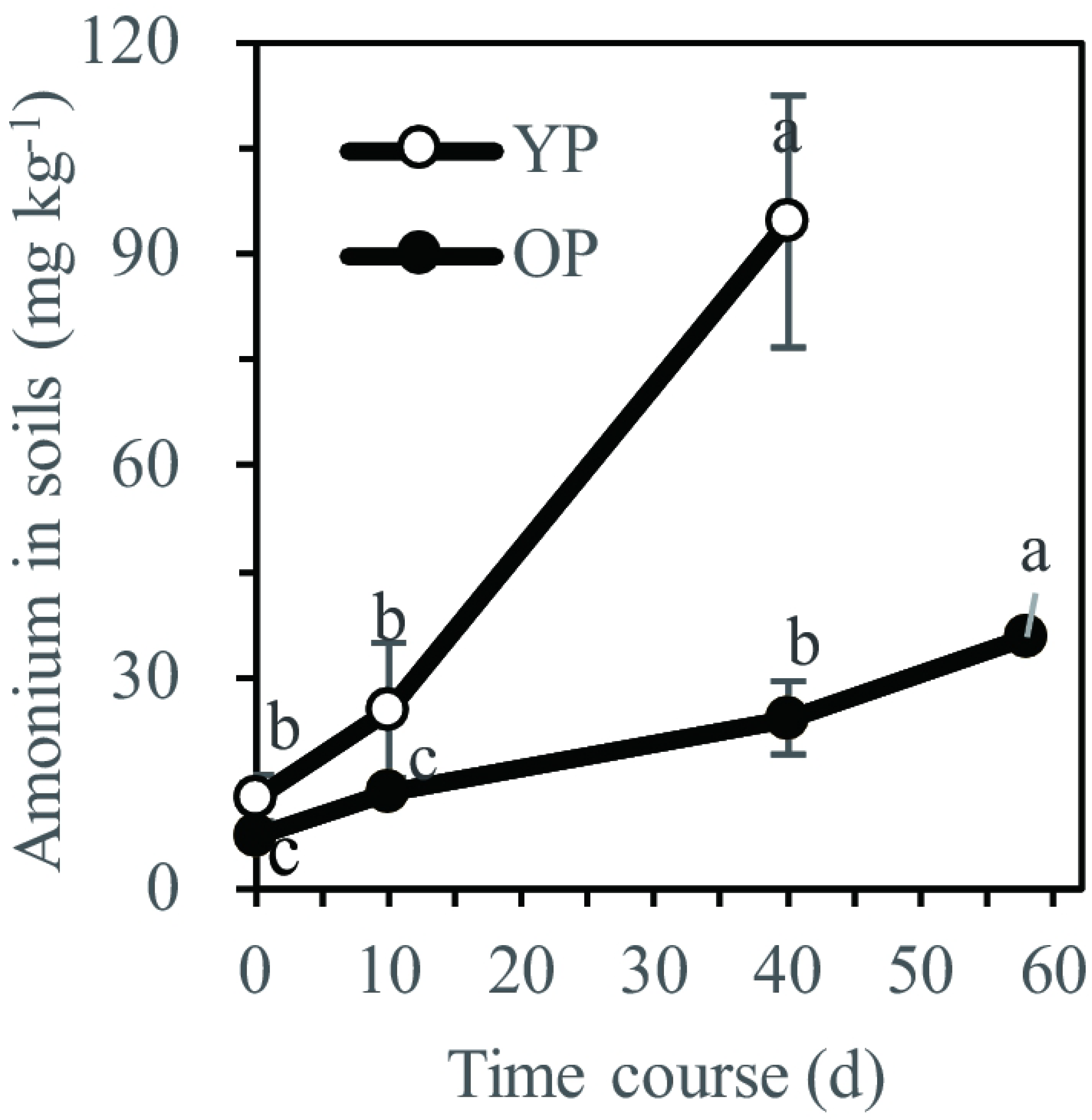
Ammonium contents in the soils. The soil samples were collected on the indicated days in 2018.

Ammonium in the medium had a positive impact on the prevention of excessive Mn uptake. The Mn content in the leaf decreased significantly after ammonium was supplied through the nutrient solution for 8 days (Fig 8). In the control plants (without ammonium application), the Mn content increased significantly with treatment time, whereas in the ammonium-treated plants, the Mn content did not decrease further after the treatment for 8 days. The Mn content in the ammonium-treated plants (98.1 mg·kg^−1^ FW) decreased by 80.2% on day 12 when compared with the Mn content (496.2 mg·kg^−1^ FW) in the control. These results clearly show the effects of ammonium in media on the suppression of excessive Mn accumulation in the plant.

**Fig 8.**
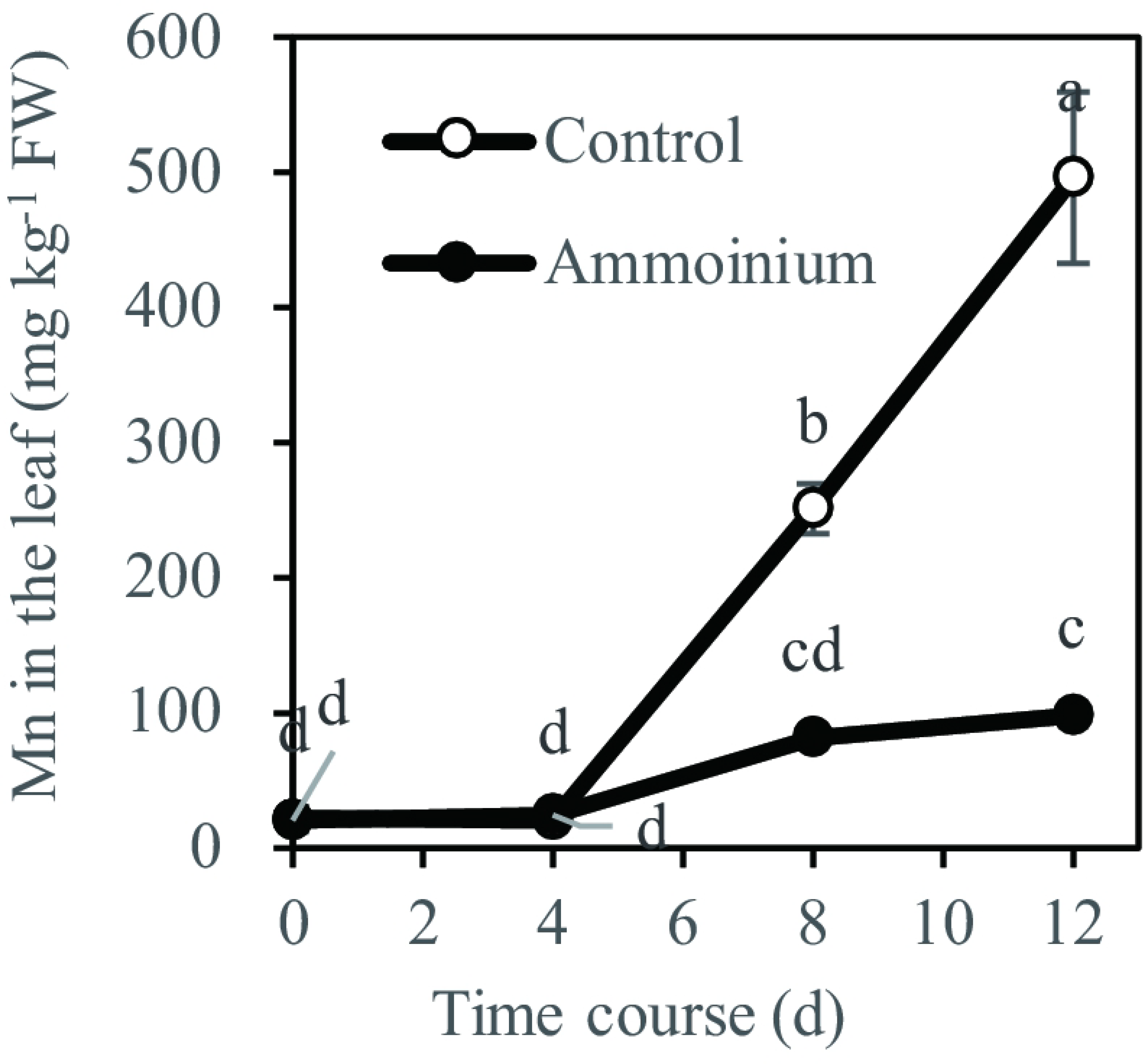
Effects of ammonium on Mn content in the first expanded leaf. Sugarcane seedlings were exposed to a nutrient solution with 0.5 mM MnCl_2_ and 0 (Control) or 1.0 mM ammonium for 0, 4, 8, and 12 days.

## Discussion

Mn is one of the most abundant elements on Earth, and excess Mn is toxic to plants. Excess Mn induces chlorosis in ratoon cane seedlings grown in acidic soils [7]. In the present study, we found that chlorotic seedlings could green in mid-summer, and this greening was associated with the downregulation of Mn content in the plants, rainfall containing Fe, and soil ammonium. To the best of our knowledge, this is the first study to demonstrate the progression of natural greening in Mn-induced chlorotic sugarcane seedlings and its association with soil and rainfall. Our findings provide the basis for an early solution for the chlorosis problem by application of Fe and/or ammonium and improvement of soil nitrogen and nutrient management in agricultural practices.

### Downregulation of Mn in plants is involved in greening of chlorotic seedlings

Self-regulation is an important mechanism for plants under environmental stress. For example, regulation of Nramp3 has been associated with Mn resistance in rice [12]. Exudation of carboxylates from roots in response to Mn has also been demonstrated as an Mn tolerance mechanism in ryegrass [13]. In the present study, we found that the symptoms of chlorosis completely disappeared in mid-summer in the ratoon cane seedlings in a natural environment without any manmade disturbances (Fig 1); the leaf SPAD values increased significantly during the progression of greening, suggesting that self-regulation mechanisms for Mn resistance act during the growth of sugarcane.

A previous study has reported the involvement of soil moisture-induced changes in available Fe in the incidence of chlorosis during season alternation. However, in the present study, no obvious differences in soil soluble Fe contents were observed during the progression of greening. In fact, the Mn content decreased in the plants during the progression of greening (Fig 3A and B). Mn content is a key determinant of the progression of chlorosis in plants. Critical toxicity content of Mn in pea, soybean, cotton, and sunflower is 300, 600, 750, and 5300 mg·kg^−1^, respectively [2]. In this study, the chlorotic seedlings showed high Mn content in their leaves, and it ranged from 266.2 mg·kg^−1^ DW to 701.0 mg·kg^−1^ DW (Fig 3A and B). The leaves of OP with 342.5 mg·kg^−1^ DW of Mn showed chlorosis, whereas the leaves of YP seedlings with 243.2 mg·kg^−1^ DW of Mn did not show visible symptoms of chlorosis (Fig 3B). All the greened seedlings, with higher leaf SPAD values ranging from 45.7 to 49.2 (Fig 2), showed leaf Mn content lower than 243.2 mg·kg^−1^ DW. These results indicate that natural greening of the chlorotic seedlings is a result of downregulation of Mn content in the seedlings.

In plants, Mn content is attributable to the integration of two dynamic processes: dry matter accumulation and Mn absorption [14]. If dry matter accumulation increases at a faster rate than Mn absorption, a decrease in Mn content is observed. This inverse relationship between growth and mineral content is termed the dilution effect [15]. Plant tolerance to Mn toxicity is, to some extent, dependent on the growth rate [2]. The incidence of Mn toxicity strongly depends on environmental factors. Sugarcane is a tropical crop with an optimum growth temperature of 30–32 °C. The atmospheric temperature increases from early summer to mid-summer and reaches 30 °C in mid-summer in the study area [7]. The chlorotic seedlings exhibited an obviously faster growth rate after mid-summer (Fig 5). A previous study has shown that plant tolerance to Mn toxicity increases with the temperature increases [16]. Fast-growing leaves form large vacuoles, thereby sequestering the potentially toxic Mn [2]. However, Mn distribution to the leaf cell sap decreased after the seedlings greened (Fig 3C). The Mn content in the plants decreased dramatically during the progression of greening (Fig 3). This decrease could be explained by the dilution effect, rather than by Mn sequestration into cell sap components such as vacuoles. The results indicate that downregulation of Mn in plants by fast growth is associated with the greening of chlorotic sugarcane seedlings.

### Rainfall contributed to greening of the chlorotic seedlings

The precipitation in the study area increased to a great extent during the progression of seedling greening (Fig S). Fe content of rainwater has been determined in previous studies [17–19], and the findings are consistent with the results of our study. Rainwater is a significant source of Fe for surface seawater and plants. More than 80% of the dissolved Fe in rain is Fe^2+^ [19]. In the present study, application of Fe-containing rainwater decreased chlorosis in the seedlings, resulting in an increase in leaf SPAD and Fe values (Fig 6). This is consistent with the results of a previous study on barley by Alam et al., who showed that application of Fe to Mn-stressed plants fully counteracted Mn-induced disorders and significantly reduced Mn contents in the plants [20]. Similarly, Mn toxicity was alleviated by Fe application to soybean [21]. Michael and Beckg reported a negative correlation between Mn^2+^ and Fe^2+^ absorption [22]. The beneficial action of Fe in reducing Mn toxicity appeared to be attributable to a mechanism that prevents the excessive absorption and subsequent accumulation of Mn in the plant [2]. The Fe content in the plants increased significantly during the progression of greening of the chlorotic seedlings, and the greened seedlings contained higher Fe content in their leaves (Fig 4). On the basis of these results, we concluded that rainfall is an important factor associated with the greening of chlorotic seedlings.

### Ammonium in the soils contributed to the greening of the chlorotic seedlings

Ammonium is a major source of inorganic N taken up by the roots of plants and the dominant N form available to plants in acidic soils [23,24]; it contributes to 79% of the total soil N solution [2]. Organic N is converted to ammonium by ammonifying soil bacteria, which are a type of mesophilic microorganisms. Ammoniation is inhibited in cold soils, and warming increases soil ammonium availability [25]. In this study, we found that the ammonium content in the soil increased significantly during the progression of greening of the chlorotic seedlings (Fig. 7), and the greening was accompanied by an increase in atmospheric temperature (Figs 2) [6]. Ammonium in the medium had a negative impact on Mn accumulation by the plants (Fig 8). N forms affect Mn toxicity [3, 4]. A previous study showed that plants grown with ammonium had less Mn content in their shoot tissue and developed no symptoms of Mn toxicity when exposed to the same Mn content [2]. Arnon reported that ammonium sources decreased Mn absorption and toxicity in barley [26]. Ammonium enhanced root-induced acidification of the rhizosphere in grass and clover [27]. Ammonium probably alleviates metal toxicity through medium pH changes and ionic competitive effects [28]. These findings suggest that soil ammonium is involved in the greening of the chlorotic sugarcane seedlings.

In conclusion, our results indicate that the chlorotic seedlings could naturally turn green, which is the result of the downregulation of Mn in the plant, supplementation of Fe via rainfall, and suppression of Mn accumulation in the plant by ammonium in the soil.

## Acknowledgments

This work was funded by the National Natural Science Foundation of China (grant No.31260497, 31660593), Guangxi Natural Science Foundation (grant No. 2016GXNSFDA380038, 2016GXNSFAA380227), Innovation Project of Guangxi Graduate Education (grant No. YCBZ2014011) and State Key Laboratory for Conservation and Utilization of Subtropical Agro-bioresources Open Foundation (OSKL201509).

## Author Contributions

Conceived and designed the experiments: XFL GZL. Performed the experiments: GZL XXW SY SJJ MMC. Wrote the paper: XFL GZL XLT.

---

**Fig S.**
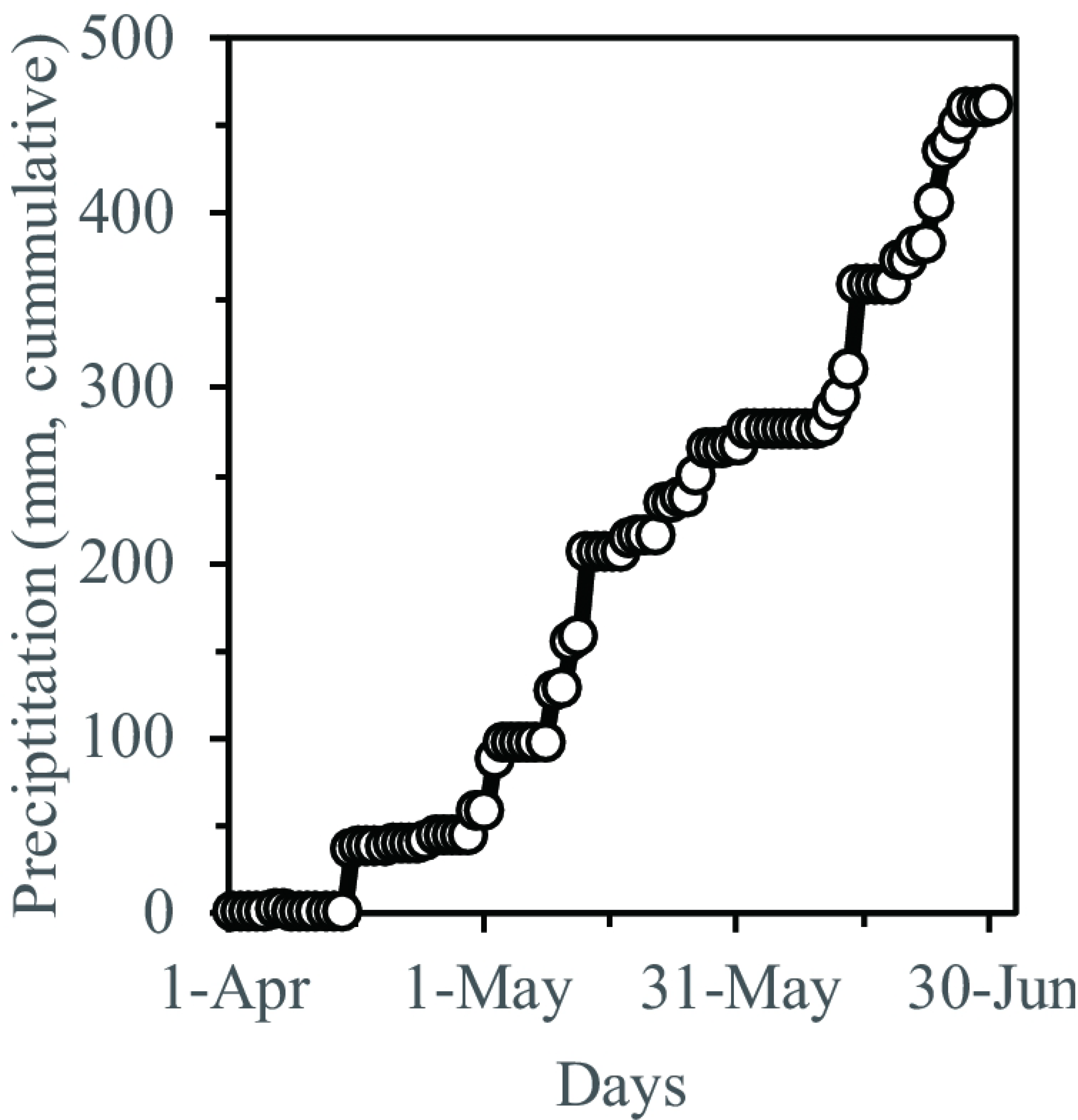
Cumulative day precipitation. Data were collected during April 1 to Jun 30, 2018.

